# Dicer- and BSC-dependent miRNAs during murine anagen development

**DOI:** 10.1101/2020.01.12.903567

**Authors:** Neda Vishlaghi, Thomas S. Lisse

**Affiliations:** The University of Miami, Department of Biology, Coral Gables, FL 33124; Sylvester Comprehensive Cancer Center, Miller School of Medicine, University of Miami, Miami, Florida 33136 USA

**Keywords:** MicroRNA, Dicer, Tarbp2, hair follicle, hair cycle, bulge stem cell, regeneration, anagen

## Abstract

MicroRNAs (miRNAs) are a major class of conserved non-coding RNAs that have a wide range of functions during development and disease. Biogenesis of canonical miRNAs depend on the cytoplasmic processing of pre-miRNAs to mature miRNAs by the Dicer endoribonuclease. Once mature miRNAs are generated, the miRNA-induced silencing complex, or miRISC, incorporates one strand of miRNAs as a template for recognizing complementary target messenger RNAs (mRNAs) to dictate post-transcriptional gene expression. Besides regulating miRNA biogenesis, Dicer is also part of miRISC to assist in activation of the complex. Dicer associates with other regulatory miRISC co-factors such as trans-activation responsive RNA-binding protein (Tarbp2) to regulate miRNA-based RNA interference. Although the functional role of miRNAs within epidermal keratinocytes have been extensively studied within embryonic and post-natal mouse skin, its contribution to the normal function of hair follicle bulge stem cells (BSCs) during post-natal hair follicle development is unknown. With this question in mind, we sought to ascertain whether Dicer-Tarpb2 plays a functional role within BSCs during induced anagen development by utilizing conditional knockout mouse models. Our findings suggest that Dicer, but not Tarbp2, functions within BSCs to regulate induced anagen (growth) development of post-natal hair follicles. These findings strengthen our understanding of miRNA-dependency within hair follicle cells to define a boundary for post-transcriptional gene regulation during anagen development.

## Background

The functional roles of microRNAs (miRNAs) within epidermal keratinocytes have been extensively studied and entail control over cell proliferation, differentiation, migration, development and epithelial-mesenchymal transitions (Mardaryev et al., 2010, Xu et al., 2011, Yi et al., 2009, Yu et al., 2010). Within embryonic skin of mice, epidermal ablation of core miRNA machinery components *Dicer, Ago1/2, Drosha* or *Dgcr8* disrupts hair follicle (HF) morphogenesis and ensuing hair formation (Andl et al., 2006, Teta et al., 2012, Yi et al., 2006, Yi et al., 2009). Additionally, overexpression studies within keratin 14-positive basal epidermal cells have also shown the inhibitory effects of miR-214 on skin and HF morphogenesis (Ahmed et al., 2014). By and large these findings depict the important role of global miRNAs toward the expansion of HF and skin cells during embryonic and neonatal development.

Only a few studies have investigated the role of miRNAs during the post-natal period within skin cells. For example, *Dicer* within keratin 5 (K5)-positive epithelial keratinocytes and/or HF epithelial stem cells (Joost S, 2019) of young mice was found to be required for post-natal HF growth and plucking-induced anagen development associated with strong phenotypes (Teta et al., 2012). Recently, miR-218-5p was shown to regulate post-natal skin and HF development by induction of the Wnt signaling pathway (Zhao et al., 2019), however the exact cell types involved remain unclear. Likewise, multiple Dicer-dependent (Lee and Doudna, 2012) miRNAs are expressed in a synchronous pattern following the post-natal hair cycle (Mardaryev et al., 2010, Zhao et al., 2019). Nonetheless, whether miRNAs within post-natal HF bulge stem cells (BSCs) - the main source of telogen to anagen transformation - play a functional role during induced anagen development in mice remains unknown. To help bridge this knowledge gap and advance the field, we conditionally ablated *Dicer* and one of its regulatory co-factors, trans-activation responsive RNA-binding protein (*Tarbp2*), specifically within BSCs during induced anagen development. Dicer-Tarbp2 regulates miRNA biogenesis (Pullagura et al., 2018), pre-miRNA cleavage specificity (Lee and Doudna, 2012) and facilitates RNA interference by association with the miRNA-induced silencing complex (miRISC) loading complex (Lee et al., 2013). Therefore, we hypothesized that global miRNAs and miRISC-mediated RNA silencing are required within BSCs during induced anagen development.

## Results and Discussion

To test our hypothesis, we performed hair depilation assays during the resting stage of the HF growth cycle (P50) using inducible *Tarbp2* and *Dicer* floxed mouse lines crossed to keratin 15 (K15) PR1Cre transgenic mice (**Figure 1A**). This system dominantly ablates *Tarbp2* and *Dicer* within outer BSCs upon RU486 treatment and not within cycling and basal interfollicular epidermal cells as previously performed (Joost S, 2019, Teta et al., 2012). By visual inspection, both control *Tarbp2*^+/+^:K15PR1Cre+ and experimental *Tarbp2*^flox/flox^:K15PR1Cre+ mice exhibited hyperpigmentation at 9 days post depilation (DPD) (**Figure 1B**). Through histological analysis, we observed the expected anagen HFs at 9 DPD among control mice (**Figure 1C**) and observed no histological nor HF growth (**Figure 1D**) abnormalities among mutant *Tarbp2*^flox/flox^:K15PR1Cre+ mice. We also applied an ultimate 3D imaging of solvent-cleared organs (uDISCO) whole-mount clearing method and observed no major difference in the extent of hair growth and hair shaft formation after depilation (**Figure 1E**). In deletion PCR studies, only RU486-treated skin samples derived from *Tarbp2*^flox/flox^:K15PR1Cre+ mice resulted in *Tarbp2* deletion (**Figure 1F**). Collectively, our results suggest that Tarbp2 regulation of miRISC within BSCs is not essential during induced anagen development of HFs.

**Figure 1.**
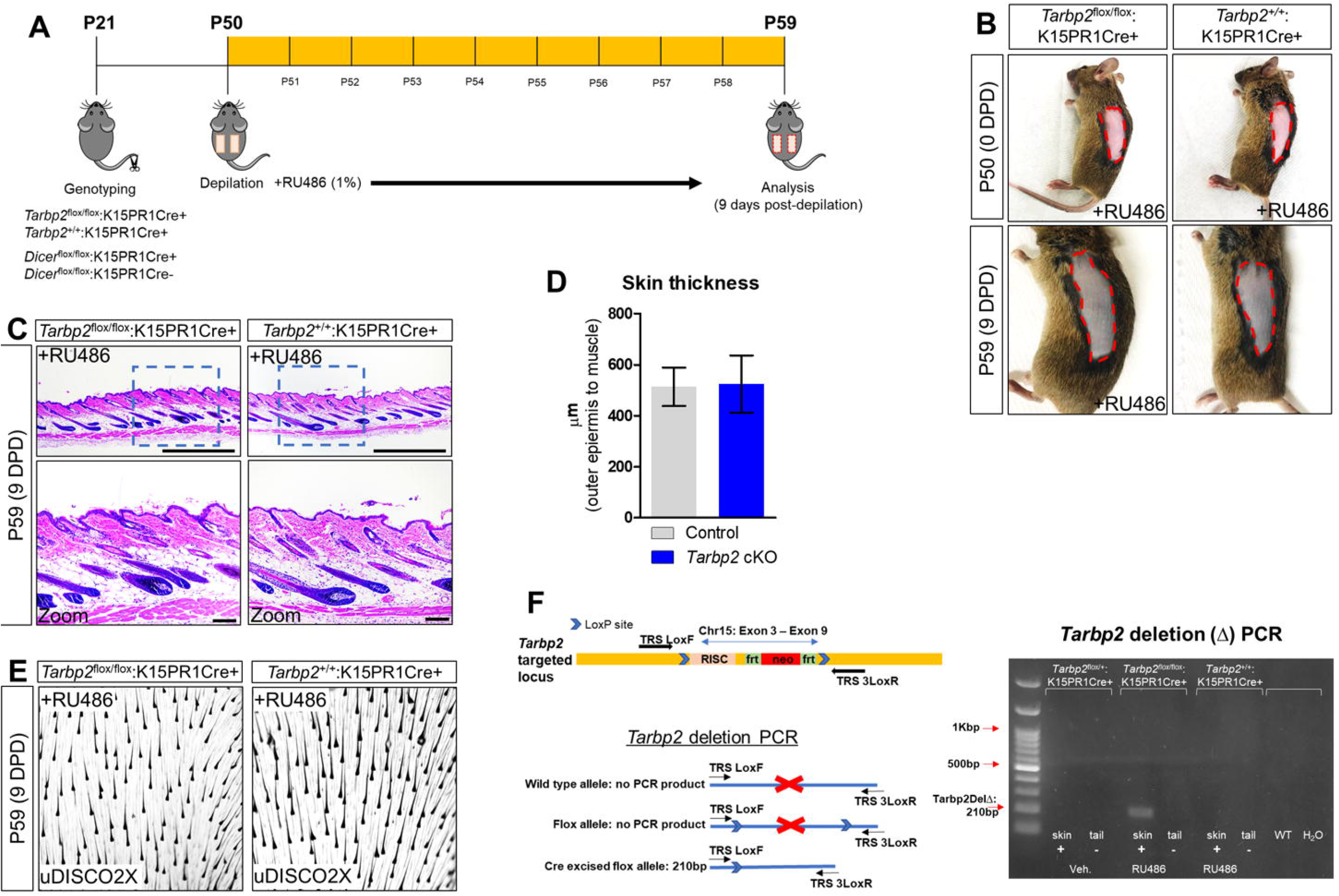
Conditional knock out of *Tarbp2* with hair follicle bulge stem cells. A) Schema outlining the genotyping and conditional knockout of *Tarbp2* and *Dicer* within bulge stem cells. B) Images of the depilated regions of control (*Tarbp2*^+/+^;K15PR1Cre+) and experimental (*Tarbp2*^flox/flox^;K15PR1Cre+) mice at 0 days post depilation (DPD) and 9 DPD. C) Histological analysis of 9 DPD skins in both control (*Tarbp2*^+/+^;K15PR1Cre+) and experimental (*Tarbp2*^flox/flox^;K15PR1Cre+) mice. Boxed region is magnified in the lower panels. Bars (top panels = 1mm; bottom panels = 100um) D) Skin thickness. Distance from the striated muscle layer to the outer epidermal layer (μm). E) Whole mount uDISCO analysis of 9DPD skins in both control (*Tarbp2*^+/+^;K15PR1Cre+) and experimental (*Tarbp2*^flox/flox^;K15PR1Cre+) mice. F) Schema of the targeting construct for generation of *Tarbp2* floxed mice (top panel). Schema of the PCR strategy to determine *Tabp2* ablation efficiency within tissue samples (bottom panel). Right panel shows the *Tabp2* deletion PCR results using RU486 pre and post-treated tissue samples.

Since Tarbp2 regulates Dicer activity (Pullagura et al., 2018), we conditionally ablated *Dicer* within BSCs by generating the *Dicer*^flox/flox^:K15PR1Cre+ mouse model and monitored anagen development after depilation (**Figure 1A**). By visual observation, both control *Dicer*^flox/flox^:K15PR1Cre- and experimental *Dicer*^flox/flox^:K15PR1Cre+ mice exhibited hyperpigmentation at 9 DPD (**Figure 2A**) and similar number of anagen HFs (**Figure 2B-D**), suggesting intact HF stem cell linage specification. However, we observed a small, yet statistically significant decrease in the thickness of skin upon *Dicer* ablation when compared to controls (**Figure 2C**), suggesting a mild delay in anagen progression (Paus et al., 1999). *Dicer* deletion PCR studies confirmed that only RU486-treated skin samples from *Dicer* mutant mice generated a deletion PCR product (**Figure 2E**). To further validate *Dicer* ablation within BSCs, we performed immunofluorescence staining for DICER using an antibody that specifically recognizes exons 22-23 of DICER (i.e. the region of *Dicer* which is excised). We observed a statistically significant (*p*≤0.001, Student *t* test) decrease in background normalized DICER immunoreactivity within the BSC compartments of mutant mice when compared to control (**Figure 2F**). Importantly, we also observed decreased DICER expression within BSC progeny of individual mutant HFs (**Figure 2G**). In control HFs, we observed DICER-positive cells in the upper ORS, combined with increased presence of DICER-positive cells within the inner root sheath (IRS) and cortex (Cx). The cellular decrease in DICER throughout mutant HFs was likely due to HF resolve to monoclonality as also shown by the K15PR1Cre+:R26R-Confetti reporter line (**Figure 2H**). Furthermore, the DICER immunostaining pattern in the control animals was consistent with recently available single cell-RNAseq data showing Dicer mRNA expression specifically within a subset of Cx/IRS as well as ORS cells within anagen HFs (Joost S, 2019) (**Figure 2I**), suggesting the cell types of miRNA dependency. Previous findings have underscored the role of miRNAs during skin development and post-natal skin maintenance by manipulating specific cell types. In summary, we show that miRNAs within K15-postive BSCs help regulate proper post-natal HF anagen development. Furthermore, we show the possibility of co-regulatory redundancy of Tarbp2 by other Dicer associated factors which may help define miRNA specificity within BSCs. Collectively, our findings demonstrate the striking cellular specificity and balance that global Dicer- and BSC-dependent miRNAs exhibit during induced anagen development.

**Figure 2.**
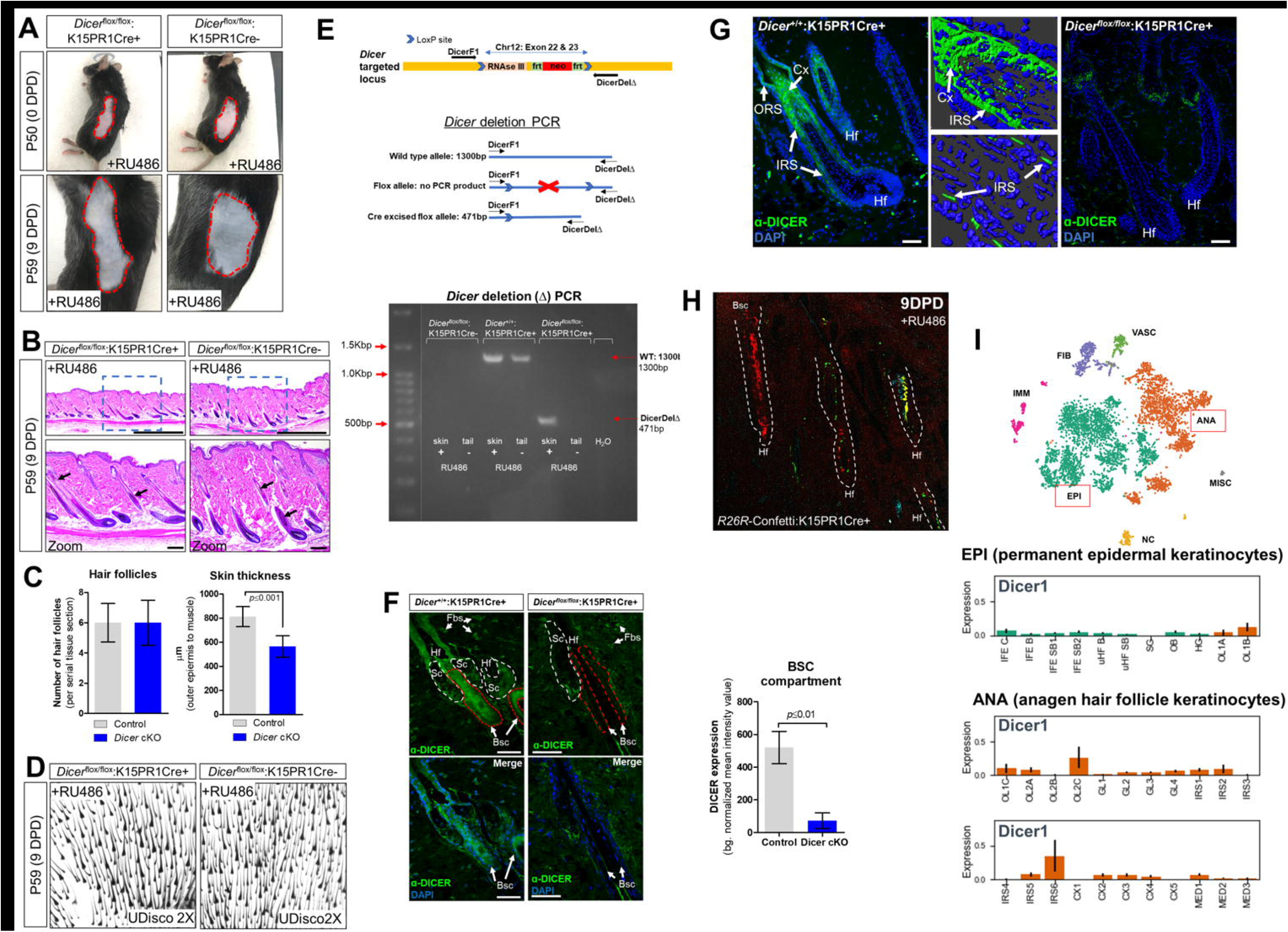
Conditional knockout of *Dicer* within hair follicle bulge stem cells. A) Images of the depilated regions of control (*Dicer*^*flox*/*flox*^;K15PR1Cre-) and experimental (*Dicer*^flox/flox^;K15PR1Cre+) mice at 0 days post depilation (DPD) and 9DPD. B) Histological analysis of 9DPD skins in both control (*Dicer*^flox/flox^;K15PR1Cre-) and experimental (*Dicer*^flox/flox^;K15PR1Cre+) mice. Boxed region is magnified in the lower panels. Bars (top panels = 1mm; bottom panels = 100um) C) Hair follicle count and skin thickness analysis. Serial skin sections were assessed from control and Dicer cKO mice (n=12;*p*≤0.001, Student’s t test). D) Whole-mount uDISCO analysis of 9DPD skins in both control (*Dicer*^flox/flox^;K15PR1Cre-) and experimental (*Dicer*^flox/flox^;K15PR1Cre+) mice. E) Schema of the targeting construct for generation of *Dicer* floxed mice. Schema of the PCR strategy to determine *Dicer* ablation efficiency within tissue samples (top panel). *Dicer* deletion PCR using RU486 pre- and post-treated tissue samples (bottom panel). F) Immunofluorescence detection of DICER within various hair follicle compartments of control (*Dicer*^+/+^;K15PR1Cre+) and experimental (*Dicer*^flox/flox^;K15PR1Cre+) skin. Upper panels depict the individual compartments and cell types that express or do not express DICER. Hf (hair follicle), Sc (sebaceous gland compartment; white dash), Fbs (fibroblasts), Bsc (bulge stem cell compartment; red dash). The lower panels highlight only the Bsc and nuclear staining with DAPI. Bars = 50um. (Right panel) Quantification of DICER expression from immunofluorescence staining. DICER expression was analyzed within bulge stem cell compartments (n=8; *p*≤0.01, Student’s t test) and presented as background normalized mean intensity values. G) Immunofluorescence detection of DICER within the entire hair follicle of control (*Dicer*^+/+^;K15PR1Cre+) and experimental (*Dicer*^flox/flox^;K15PR1Cre+) skin. Right panels depict 3D-rendered models of DICER expression in control hair follicles. ORS (outer root sheath), IRS (inner root sheath), Cx (cortex). Bars = 50um. H) K15 BSC lineage tracing using the *R26R*-Confetti:K15PR1Cre+ mouse line. I) Dicer mRNA expression at the single cell level during hair growth (anagen) and rest (telogen) in full-thickness skin. Permanent epidermis keratinocytes (shown in green) contain subpopulations of resting BSCs and epidermal keratinocytes. The anagen hair follicle keratinocytes (shown in orange) contain subpopulations of cells undergoing growth and differentiation. EPI (permanent <u>epi</u>dermal keratinocytes), ANA (anagen hair follicle keratinocytes). EPI cells consist of interfollicular epidermis (IFE) basal cycling (C), basal (B) and suprabasal (SB) cells, upper hair follicle (uHF) basal (B) and suprabasal (SB) cells, sebaceous glands (SG), outer bulge (OB), and hair germ (HG) cells. ANA cells consist of outer layer (OL) cells, germinative layer (GL) cells, inner root sheath (IRS) cells, cortex/cuticle (CX) cells, and medulla (MED) cells. Data derived from: (Joost S, 2019)

## Data Availability

The data supporting the results reported in this article will be provided upon reasonable request.

## Author Contribution

N.V. was involved in the formal investigation, methodology, analysis of the data, and the review and editing of the manuscript. T.S.L. was involved in the conceptualization, project administration, formal investigation, methodology, analysis of the data, and writing of the original draft of the manuscript.

## Acknowledgement

We especially thank Legnay Fernandez (U. Miami) and Kathryn Helen Forcone (U. Miami) for their critical comments on this manuscript. We especially thank the Animal Care and Veterinary staff at the Neuroscience Annex (U. Miami) for their animal support.

## Materials and Methods

### Mouse Models and Breeding

*Dicer*- Null conditional (*Dicer*1^*tm1Bdh*^/J) mice (referred to as *Dicer^floxed^*; JAX stock no. 0063661) and *Tarbp2*-Null conditional (*Tarbp2^tm1.1Dzw^*) mice (referred to as *Tarbp2*^*Floxed*^;MGI: 5645258) were received from the Jackson laboratory. The Tg(Krt1-15-cre/PGR)22Cot line (referred to as *K15*-PR1Cre; JAX stock no. 005249) was on a mixed C57BL/6 and SJL background also obtained from Jackson Laboratory and was used for generation of RU486-induced, Cre-mediated gene deletions within BSCs of hair follicles (HFs). *Dicer^flox/flox^* (*Dicer^fl/fl^*) and *Tarbp2^flox/flox^* (*Tarbp2^fl/fl^*) mice were bred with K15-Cre line to achieve cell-specific knockout of targeted regions within BSCs of HFs. Lineage tracing was performed using the *Gt(ROSA)26Sor^tm1(CAG-Brainbow2.1)Cle/J^* mice obtained from the Jackson Laboratory (R26R-Confetti; JAX stock no. 013731). All animals were maintained in a 12-h light (6am to 6pm) and 12-h dark cycle vivarium in the Research Animal Facility at the University of Miami. Animals were provided acidified tap water through filter bottle and irradiated pelleted 2018 Teklad global 18% protein rodent diet from ENVIGO. In timed pregnancy crosses, pups were dated based on the presence of vaginal plugs and by noting the delivery of newly born pups. All animal studies were approved by the University of Miami Institutional Animal Care and Use Committee (IACUC) protocols.

### Genotyping and recombination detection

All genotyping proceeded by using tail tip excision/partial amputation under the age of 21 days. Dicer floxed allele was genotyped by using primers DicerF1 (CCTGACAGTGACGGTCCAAAG) and DicerR1 (CATGACTCTTCAACTCAAACT). PCR product for genotyping PCR was 420-bp band for Dicer and a 351-bp band for wild type allele. The deletion was genotyped by using primers DicerF1 and DicerDel (CCTGAGCAAGGCAAGTCATTC). PCR product for deletion PCR was a 471-bp band for deletion and a 1300-bp band for the wild type allele. To assay floxed vs. wild type allele in Tarbp2 animals, we used TRS-loxF (CAGAAGCACAGCAGGAACAA) and TRS-loxR (CGTGATATGCACAGCCCACT) primers. PCR product for genotyping PCR was 180-bp band for the floxed and a 130-bp band for the wild type allele. Deletion allele was detected by using primers TRS-loxF and TRS-3loxR (CAAAACCACTTCCCCATGTT). All mice genotyping was based on established protocols listed on the JAX website for each stock animal. PCR program was run according to recommendations: 95°C for 5 minutes, and for 35 cycles 95°C for 15 seconds, 60°C for 15 seconds and 72°C for 10 seconds for Dicer genotyping and deletion PCR; 94°C for 3 minutes, for 35 cycles 94°C for 30 seconds, 61.7°C (59°C annealing for delete allele) for 30 seconds and 72°C for 20 seconds for Tarbp2 PCR.

### Conditional knockout studies and hair depilation assay

Under general anesthesia (inhalation isoflurane), 50-day-old animals were subjected to hair depilation of two separate left and right side of the dorsal area by using Wax Strips. All procedures were performed using sterile instruments and aseptic conditions. Mice received topical daily treatments of 1% RU486 in acetone, or acetone only (vehicle), over the depilated area for nine days. Nine days after depilation, animals were sacrificed, and skin samples were collected for analysis.

### Ultimate DISCO (uDISCO) whole mount passive clearing technique for skin

To visualize thick skin tissue and hair morphology, we applied the organic solvent-based uDISCO clearing method. Greater than 3mm-by-3mm skin tissues were fixed in 4% PFA at 4°C overnight. Samples were washed with 0.1M PBS and subjected to a tert-butanol series (70-100%; Sigma-Aldrich, 36053) for gradient dehydration for 2-12 hours. Next, dichloromethane (Sigma-Aldrich, 270997) was replaced as a pure solution for the delipidation step for 1 hour at room temperature. A refractive index matching solution was prepared by mixing BABB (benzyl alcohol + benzyl benzoate 1:2; Sigma-Aldrich, 24122 and W213802) and DPE (diphenyl ether) (Sigma-Aldrich, 240834) at a BABB:DPE ratio of 10:1 (vol/vol). The samples were reacted with the BABB:DPE mixture until they became optically transparent.

### Histological, histomorphometric and immunofluorescence analysis

Skin tissues (1×1 cm^2^) were dissected from mice and incubated 2-4 hours in 4% PFA in PBS (pH 7.4) at 4°C and transferred to 70% OH, before embedding in paraffin wax. 7μm-sections were stained with hematoxylin and eosin (H&E) by The University of Miami Histology Core Services. For immunofluorescence studies, skin tissues (1×1 cm^2^) were dissected from mice and incubated 2-4 hours in 4% PFA in PBS (pH 7.4) at 4°C and subjected to graded sucrose treatments (15-30%) for cryoprotection. These tissues were embedded face down along the midline in OCT embedding medium (Histolab). Cryosections were made using a Leica CM 1850 Cryostat. 10μm-sections were incubated with 0.4% Triton X-100 in PBS for 30 minutes at RT to reduce background staining. Tissue sections were directly blocked in PBS containing 5% normal horse serum for 30 minutes at RT. Then incubated with endogenous Mouse IgG blocking solution 1:10 in PBS (Unconjugated AffiniPure Fab Fragment Goat Anti-mouse IgG(H+L); Jackson ImmuneResearch Labs, 115-007-003) for 1 hour at RT. Sections were incubated with primary antibodies at optimal dilution for 30 minutes at RT, antibodies used in this study included Dicer (BioLegend. 820201). Following 3x washes for 2 minutes each in PBS-Tween-20, the sections were incubated at room temperature for 20 minutes with corresponding species-specific secondary antibodies (Alexa series, Life Technologies). Following 3x washes in PBS-Tween-20, sections were mounted with Vectashield medium containing DAPI (Vector Laboratories) for nuclei staining. Immunofluorescence microscopy was performed using a Zeiss Observer 7 ApoTome2 unit. Images were captured from BSCs compartment for DICER expression and normalized to acellular dermal regions. Means of expression intensity of individual follicles were compared between control and mutant animals (n=8). Imaris (Bitplane) was used to generate 3D rendered models of Dicer expression. For hair follicle counts (i.e. hair follicles per 2.5mm tissue; n=12) and skin thickness measurements (i.e. distance from panniculus carnosus (muscle) to the outer epidermis; n=12), serial skin sections were analyzed between control and cKO mice. Two-tailed unpaired T tests were performed between control and cKO data sets using Prism (GraphPad) where the *p* value summaries were depicted as \*\*\**p* ≤ 0.001 \*\**p* ≤ 0.01, and \**p* ≤ 0.05.

